# How many, and when? Optimising targeted gene flow for a step change in the environment

**DOI:** 10.1101/341339

**Authors:** E. Kelly, BL. Phillips

## Abstract

Targeted gene flow is an emerging conservation strategy that involves translocating individuals with particular traits to places where they are of benefit, thereby increasing a population’s evolutionary resilience. While the idea can work in theory, questions remain as to how best to implement it. Here, we vary timing of introduction and size of the introduced cohort to maximise our objective – survival of the recipient population’s genome. We demonstrate our approach using the northern quoll, an Australian marsupial predator threatened by the toxic cane toad. We highlight a general trade-off between maintaining a local genome and reducing population extinction risk, but show that key management levers can optimise this so that 100% of the population’s genome is preserved. In our case, any action was better than not acting at all (even with strong outbreeding depression), but the size of the benefit was sensitive to timing and size of the introduction.

## Introduction

Rapid environmental change is causing declines in biodiversity across the globe (Barnosky *et al.* 2011), and as threats become harder to mitigate, threatened species must adapt to survive (Hoffmann & Sgrò 2014). Existing genetic variation in relevant traits coupled with strong selection imposed by a threatening process may allow some populations to rapidly adapt; staving off extinction through evolutionary rescue (Bolnick *et al.* 2011; Sih *et al.* 2011). Unfortunately, for many threatening processes, beneficial traits are either locally absent or at very low frequencies, and this makes extinction more likely than evolutionary rescue (Gomulkiewicz & Holt 1995; Frankham 2015). Targeted gene flow has emerged as a conservation strategy for helping promote these favourable traits in threatened populations (Kelly & Phillips 2016). This strategy involves translocating individuals with key traits to areas where the traits would have a conservation benefit.

Although targeted gene flow has yet to be implemented and robustly assessed, it could provide benefits for populations facing a wide range of threats – such as disease, invasive species, or climate change. It can enhance the capacity of these populations to adapt to predictable and imminent changes in their environment, and so skew outcomes towards evolutionary rescue rather than extinction (Kelly & Phillips 2016). By introducing specific individuals selected for their traits, we can increase overall genetic diversity in threatened populations as well as the frequency of key traits (Tallmon *et al.* 2004; Whiteley *et al.* 2015). This leads to faster adaption to the threat, and therefore an increased chance of evolutionary rescue. By increasing the chance of evolutionary rescue, we also promote the persistence of the local genome, and so conserve genetic diversity at the species level: we aim to manipulate populations so that they are not only locally adapted, but also carry genes that allow them to survive the current threat (Kelly & Phillips 2016).

Here, we address questions on the implementation and risks of targeted gene flow using a population viability analysis that incorporates evolution. Conservation managers considering any adaptive relocation must decide on the optimal timing for their translocation, as well as the composition of the introduced cohort (McDonald-Madden *et al.* 2011). Targeted gene flow is no different, with timing and number of introductees likely to have large effects on the benefits for conservation. Selection, hybridisation and outbreeding/inbreeding depression also have strong influences on population persistence, but are rarely included in models of population viability (Pierson *et al.* 2015). For those that do, very few look at the potential impact of outbreeding depression, and none have yet assessed the potential use of targeted gene flow. For any conservation action, it is wise to assess the action prior to implementation – particularly to determine the best way to implement the strategy, and to assess possible negative effects (Coulson *et al.* 2001). We addressed these key aspects of conservation decision making (timing and number of introductees) in our model. We also examined how a key uncertainty (the severity of outbreeding depression) changes outcomes.

We seek a strategy that maximises expected return for conservation, but to do this requires a clear statement of our management objective (Regan *et al.* 2005). In our case we would like to keep our recipient population extant, but we would like to do so without replacing the local genome. Replacing the local genome with a genome from elsewhere is equivalent to a reintroduction, but one of the main potential benefits of targeted gene flow is the possibility that we preserve local genetic diversity. A sensible objective, then, is analogous to a gambler’s expected return: the probability of winning, multiplied by the payout. In our case the payout is the proportion of the recipient population’s genome still extant, (calculated as mean proportion of recipient genome within each individual; *r_I_*), and our probability of winning it, 1 – *p_e_*, where *p_e_* is the extinction probability. Thus, our objective is to maximise the expected return:

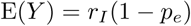

We then chose a case study: the endangered carnivorous marsupial, the northern quoll (*Dasyurus hallucatus*), a potential candidate for targeted gene flow. Northern quolls are threatened by the invasion of the toxic cane toad (*Rhinella marina*) because quolls unknowingly eat the toxic toads and are fatally poisoned (Tingley *et al.* 2017). The great majority of quoll populations (more than 95%) go extinct after cane toad arrival (EPBC 1999), which represents a large step-change in their environment. Cane toads continue to spread westward, and will eventually inhabit the quolls’ entire range (Tingley *et al.* 2017). Evidence suggests a small number of scattered populations have adapted to the threat (Kelly & Phillips 2017, 2018) by evolving to avoid toads as prey. Juvenile northern quolls born in a toad-free environment will avoid cane toads if they have at least one parent from one of these toad-smart populations (Kelly & Phillips 2018), indicating that toad-smart behaviour has a heritable basis. It is this “toad-smart” behavioural trait that we can make use of targeted gene flow. Here, we use northern quolls as a model for optimising targeted gene flow following a step change in the environment (in this case, the arrival of cane toads). We use an individual based model and population viability analysis, varying key management objectives (timing of the introduction and size of the introduced cohort) to maximise the expected return for conservation.

## Methods

We developed a discrete-time individual based population model. The demographic aspects of the model are tailored to northern quolls, with age- and sex-specific annual survival and fecundity rates. Demographic stochasticity (stochastic survival and reproduction) is introduced at the individual level, and each individual expresses a trait determining the individual’s probability of being killed by a cane toad. Toad-driven mortality was imposed only in the first year juvenile stage. Animals surviving this event were considered to have learned to avoid toads thereafter.

### Demography

Within time intervals, reproduction (including pre-weaning survival of babies) is followed by survival of juveniles and adults. Baseline survival probabilities and fecundity were inferred for this model using the average of values drawn from the literature (Table S1).

### Female fecundity and survival of babies

Fecundity of females was considered density dependent. We considered that all females have the capacity to produce 8 babies, but that the survival of these babies to weaning is density dependent: declining from a base survival rate with an increasing density of adult females in the population. Male density was ignored because almost all adult males die before young quolls are weaned (Dickman & Braithwaite 1992). Thus, the expected number of weaned offspring for each female is 8*s_b_*, where *s_b_* is the probability that each baby survives to weaning. This survival probability is dependent on the density of adult females in the population, *n* such that:

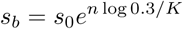

*s*_0_ is the base survival rate in the absence of density effects. Here *K* is the density of females at which survival probability of babies equals 0.3*s*_0_. The constant, 0.3, is chosen as approximately the value of *s_b_* at which the population stops growing when *s*_0_ = 1. While this process gives us an expected number of weaned offspring, for each female, the realised number of weaned offspring is stochastic; determined, for each female, as a draw from a binomial distribution Binom(*s_b_*, 8).

### Survival of juveniles and adults

We used data from previous mark-recapture studies to estimate yearly survival probabilities of male and female quolls (Begg 1981; Schmitt *et al.* 1989; Braithwaite & Griffiths 1994; Oakwood 2000). We used raw published data that indicated age and sex of the mentioned quolls, and that followed the individuals for at least one year, as this meant we could estimate annual survival (Table S1). We took the unweighted means of these data to get the survival probabilities that were used in our model. Unfortunately, there was no way of accounting for survey effort or detection probability, as the papers we used did not report these details. However, our estimates were extremely close to those from Cremona et al(Cremona *et al.* 2017), which used their own mark-recapture study and estimated annual survival incorporating recapture probability. Therefore, we believe our parameter estimates are the best that can be determined, given the available data.

Juvenile survival probability (*s*_0_) was set to 0.38 based on estimates from Oakwood (2000) of the survival of juveniles between when they were first denned to when they first became trappable. Male quolls mature within a year and, with very high probability, die before their second year. In our model, this was captured with two parameters: *s_m_*_1_ = 0.042, *s_m_*_2_ = 0. Female quolls also mature within a year, but have a greater chance to survive through to a second reproductive season. In our model, this was captured with three parameters: *s_f_*_1_ = 0.234, *s_f_*_2_ = 0.03, and *s_f_*_3_ = 0. In all cases, the realised survival of an individual was treated as a draw from a Bernoulli distribution with the specified survival probability.

### Sexual reproduction

The model tracks males and females separately. Loci are assumed not linked and so recombination rate between loci is 0.5. Mate choice is random and males can mate with multiple females, but multiple paternity (within a litter) is not allowed. Each offspring from each pair inherits a genotype determined by the fusion of gametes from the sire and dam. In all cases, a gamete is the result of random segregation of the parent’s genotype into haploid form.

### Evolutionary dynamics

Each individual expresses a continuous trait, *A,* determining whether or not the individual will eat a toad (and so die), or not. The trait is determined by the animal’s genotype, and also by environmental variation according to the underlying mechanisms of a simple quantitative genetic model in which the total phenotypic variance is the sum of genetic and environmental contributions: *V_T_* = *V_G_* + *V_E_.* Within all our simulations, the environmental variation imposed on the trait remains constant.

### The genotype and expected trait value

The animal’s genotype consists of a number of diploid, biallelic loci. A subset of these loci, *n_i_* are involved in incompatibility and another subset *n_n_* were neutral and used to track the recipient genome (see below). The remaining *n_p_* loci contribute to the organism’s phenotype. Each phenotype-influencing locus has an equal additive effect on the individual’s expected trait value, E(*A*). Two alleles are possible at each locus, with alleles having an additive effect size of either 0, or *e,* where *e* represents an increment towards being more toad-smart.

The effect size, *e* is calculated as a function of the environmental variance, and the heritability, and is chosen such that the stated heritability is achieved given the stated environmental variance and number of loci. Under a simple quantitative genetic model,

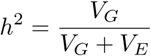

or equivalently,

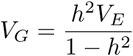

Under a normal approximation to the binomial distribution, the expected genetic variance is,

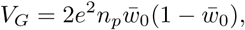

where 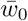 is the frequency of alleles with non-zero effect sizes. Thus, our effect size can be calculated as:

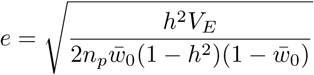

An individual’s genotypic value (its expected phenotype) is then:

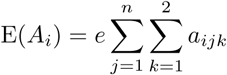

where *a_jk_* references the allelic value (either 0 or 1) at allele *k* of locus *j. i* indexes the individual.

The heritability of the trait (*h*^2^) was set to 0.2, based on the estimate of behavioural trait heritability given by (Roff 2012). Sensitivity analysis investigating the effect of *h*^2^ is presented in supplementary material (Fig. S3 & S4). We assume that the population initially has low mean fitness, 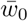, with regard to toads, and that alleles conferring toad-smart behaviour are rare in the population. We set 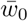 to be the initial frequency of toad-smart alleles, and this was determined by running toad introduction simulations with different values of 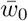 to get the model to mimic the observed population extinction rate (with no management intervention(EPBC 1999)) of 0.95 after toads arrive. 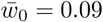 was the first value of 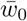 that gave a probability of extinction above 0.95, with all other parameters set constant (Fig. S1).

### The phenotype

An individual’s realised phenotypic value, *A_i_* incorporates environmental variation on the expected trait value, and is determined stochastically, as a draw from the normal distribution.

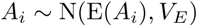

### Selection

The model implements threshold selection in which all individuals with trait values for *A_i_* ≤ *A\**, will be killed by toads, while all individuals with *A_i_* > *A\** will survive. The selection threshold, *A\**, is defined as the 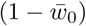th quantile of the initial expected phenotype distribution, and so is determined as a function of the environmental variance and the heritability at initialisation. The phenotype distribution is approximated as,

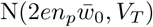

Where *V_T_* is the total phenotypic variation, *V_G_* + *V_E_.*

Given that our initial mean fitness is also the frequency of toad-smart alleles in the population, simulations that are initialised with a large number of individuals will tend to generate populations in which there is sufficient standing variation such that selection can, in principle, achieve a mean toad-smart fitness of 1. That is, across a large starting population, a toad-smart allele will tend to be present at every locus, though it might only be represented at that locus in a few individuals in the population. As is likely the case in reality (small population sizes and short time spans), selection acts on this standing variation in the population; we do not consider mutation important (???).

### Loci involved with incompatibility

To allow us to incorporate outbreeding depression, each individual also carries *n_i_* loci involved with incompatibility. These loci carry fixed differences between recipient (=0) and introduced (=1) populations.

The loci involved with incompatibility were used to implement outbreeding depression using the model of two-locus incompatibilities developed by (Turelli & Orr 2000). This model works on the theory that the sterility and inviability of hybrids can be explained by between-locus “Dobzhansky-Muller” incompatibilities. The Turelli and Orr model includes three types of incompatibilities: those between heterozygous loci (*H*_0_), those between a heterozygous and a homozygous (or hemizygous) locus (*H*_1_), and those between homozygous loci (*H*_2_). Using this model, we determine the “hybrid breakdown score” E(*S*) of each individual based on the composition of alleles in their set of loci involved with incompatibility (proportion of loci that are homozygous from population 1 (*p*_1_), the proportion that are homozygous from population 2 (*p*_2_), and the proportion that are heterozygous for material from the two populations (*p_H_*)). Following Turelli and Orr, the hybrid breakdown score is given as,

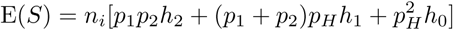

where *n_i_* is the number of loci that are contributing to incompatibilities. We then used a simple negative exponential function to link hybrid breakdown score to fitness,

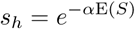

where *α* is a constant value, and *s_h_* is the probability of survival from outbreeding depression. When outbreeding depression is activated in the model, all baseline survival probabilities are multiplied by individual *s_h_* values, and survival is then determined by a draw from a Bernoulli distribution with the resultant survival probability.

We set the parameters of *n_i_* = 10 and used simple dosage ratios for the different classes of incompatibilities: (*H*_1_) = 0.5, (*H*_2_) = 1 and (*H*_0_) = 0.25, to generate the hybrid breakdown score. We then varied the value of *α* in the fitness function to manipulate the strength of outbreeding depression.

### Neutral loci

To track the fate of the introduced and recipient genomes, we also initialised *n_n_* neutral loci, also carrying fixed differences between recipient (=0) and introduced (=1) populations. We used these neutral loci to track the proportion of the recipient population’s genome remaining. This was measured in two different ways — the first (*r_I_*) was calculated as the proportion of recipient population alleles within each individual, averaged over all individuals in the population. This was used, along with probability of extinction (*p_e_*) to calculate the expected return of our management objective (E(*Y*): the proportion of the local genome surviving) for each scenario. Therefore, maximum expected return equated to the population with the highest proportion of recipient genome likely to survive the toad invasion (calculated by E(*Y*) = *r_I_*(1 – *p_e_*)).

*r_I_*, however, underestimates the true proportion of the recipient genome still extant in the population, because the original genome will be scattered across individuals within the population. Therefore, we also calculated *r_P_*, the proportion of loci which retained recipient alleles across the population. This gives us the true estimate of recipient genome retention, because it counts alleles even when they are at very low frequency (e.g., represented in only a single individual). While *r_P_* is the true proportion of the recipient genome still extant, it will likely decline over time due to loss of rare alleles through drift. Because of this, we ran our optimisation using the more conservative metric, *r_I_*, and make a comparison of the two metrics in the supplementary material (Fig. S2).

### Scenarios

All our scenarios begin with an initially poorly adapted quoll population of 1000 individuals — our “recipient population”. The recipient population is initiated with 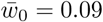 (initial frequency of toad-smart alleles in a toad-naïve population giving observed population extinction rates(EPBC 1999)) and number of loci was set to 30 (*n_i_* = 10 relating to incompatibility, *n_p_* = 10 relating to the trait and *n_i_* = 10 neutral loci). Carrying capacity (*K*) of breeding females was set to 1000. The recipient population was allowed to grow for 30 years before the environment was changed to simulate the arrival of toads. The arrival of toads was treated as a step change in the environment, happening at the beginning of a generation. This sudden change reflects the reality of toad arrival (Phillips *et al.* 2007).

In each scenario we recorded whether the population went extinct or not over 50 years following toad arrival. For populations that survived we also calculated recipient population’s genome remaining, and using these measures, calculated the maximum expected return — the population with the highest proportion of recipient genome likely to survive the toad invasion.

### Targeted gene flow with no outbreeding depression

To simulate targeted gene flow we introduced a number of toad-smart quolls to our recipient population prior to reproduction in a specified year. Our “source population” was a toad-smart population, initialized with 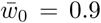 (i.e. toad-smart genotype), age = 1, and sex randomly allocated. We ran a number of scenarios to determine the effect of varying the timing of introduction and the number of introductees. We examined a range of introductee number, from 0-50 individuals (in increments of 5 individuals); and a range of introduction times, from ten years prior to toad arrival to ten years after toad arrival (in 1 year increments). We ran 100 simulations for each scenario to estimate extinction probability, *p_e_.*

### Targeted gene flow with outbreeding depression

We then ran the same simulation as above but added in varying strengths of incompatibility-driven outbreeding depression. We set outbreeding depression to have a low and high impact on hybrid individuals by reducing the fitness of F1 hybrids by 10% and 50% of baseline fitness. This was achieved by changing the value of *α* that converts hybrid breakdown score into fitness (10% reduction in F1 hybrids *α*=0.04; 50% reduction in F1 hybrids *α*=0.28).

## Results

### Targeted gene flow with no outbreeding depression

We found that the success of targeted gene flow was strongly influenced by the timing of the introduction, and the number of individuals introduced (Fig. 1). Our management objective (*Y*) was optimised when a larger number of individuals (25+) were introduced in the years immediately preceding toad arrival (Fig. 2).

**Fig. 1.**
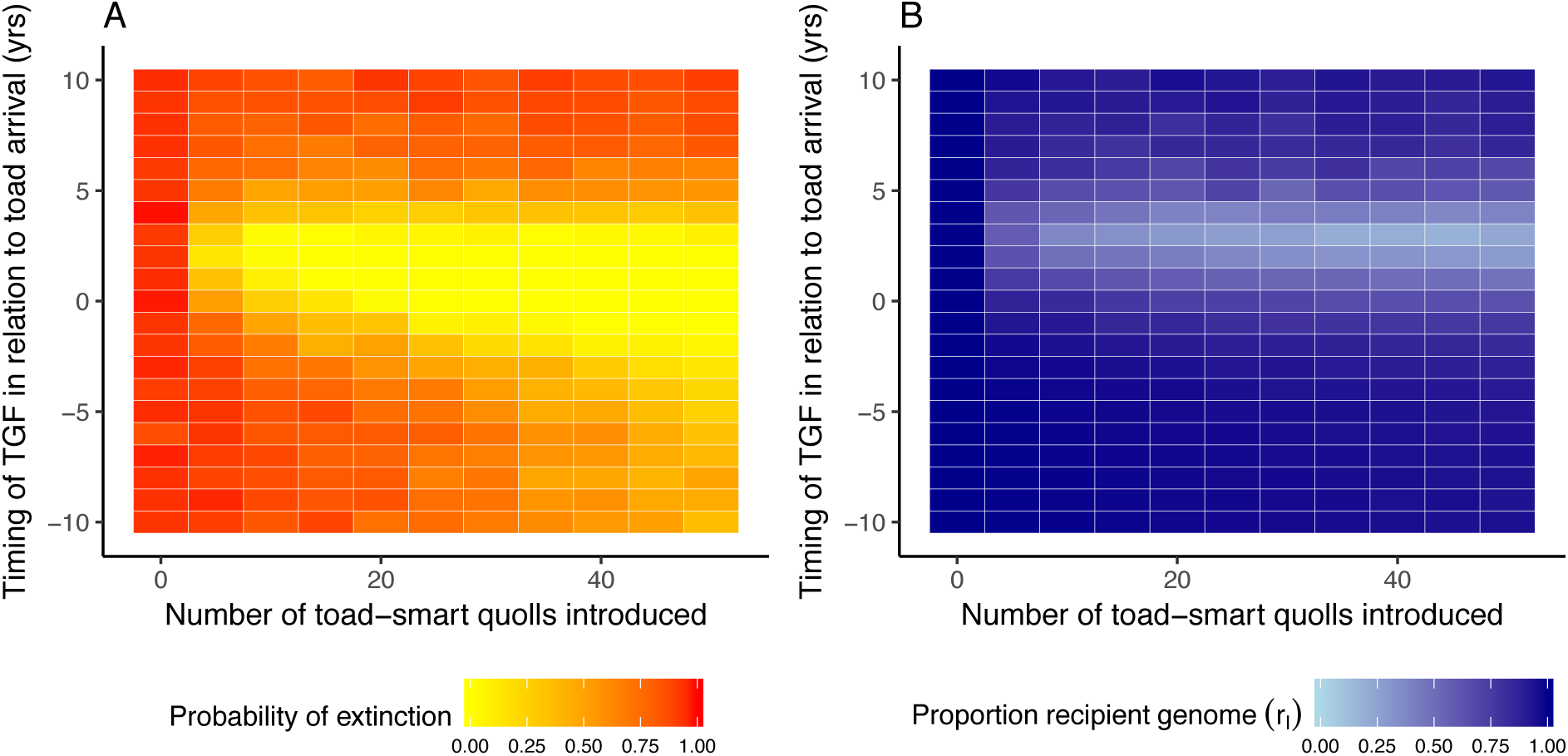
A: The probability of extinction of a northern quoll population (*p_e_*; red = high chance of extinction) for varying implementations of targeted gene flow. B: The proportion of recipient population genome (*r_I_*; dark blue = recipient genome) in eventual population after varying implementations of targeted gene flow.

**Fig. 2.**
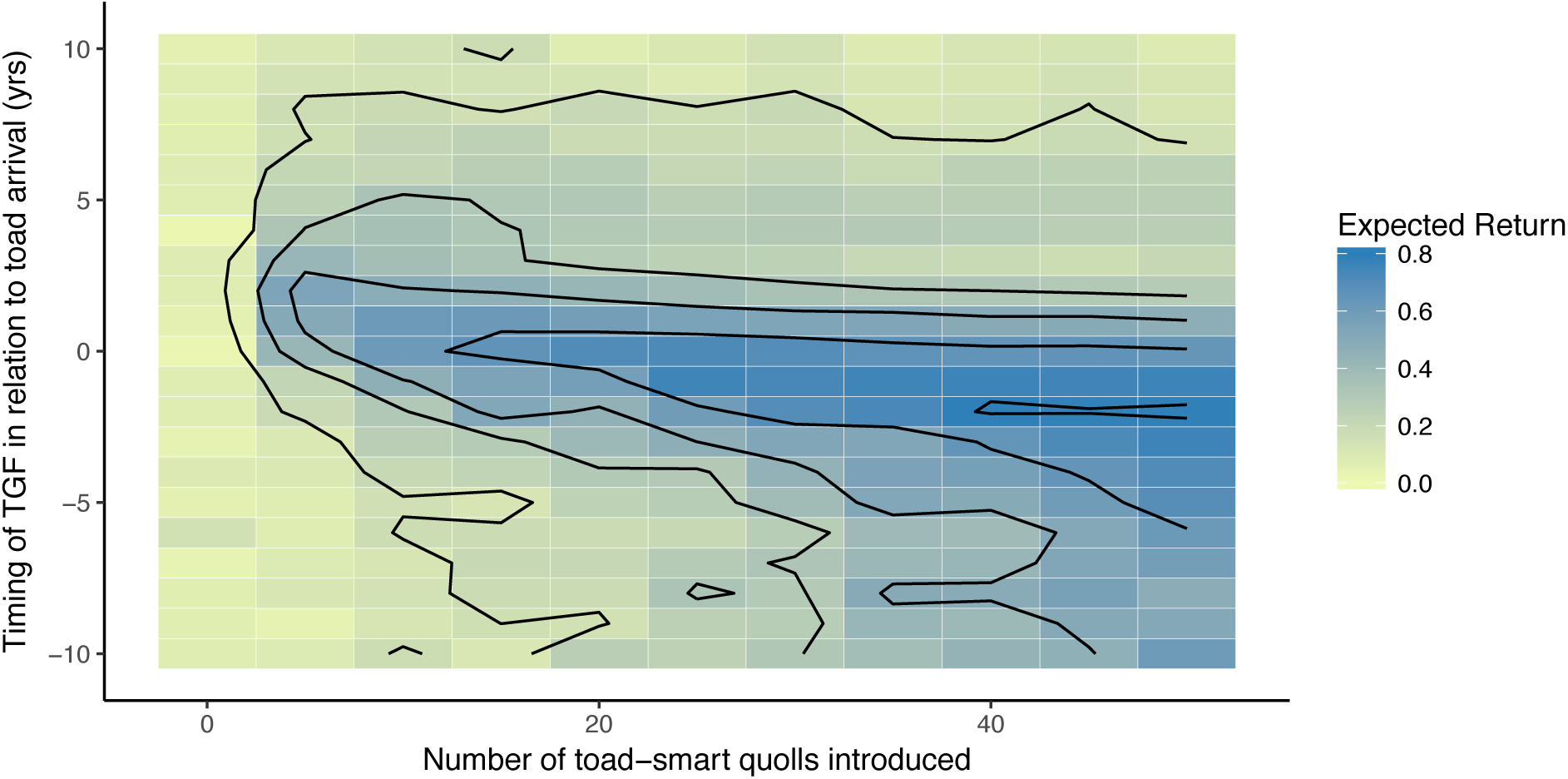
Expected return of the recipient genome (calculated by E(*Y*) = *r*(1 – *p_e_*) using probability of extinction (*p_e_*) and proportion of recipient genome (*r_I_*)).

### Targeted gene flow with outbreeding depression

Generally outbreeding depression had a negative impact on the success of targeted gene flow. 10% reduction in fitness produced relatively similar results to no outbreeding depression, however a 50% reduction in fitness increased the probability of extinction and decreased the proportion of recipient population genes, reducing the proportion of scenarios with a higher value of *Y* (Fig. 3).

**Fig. 3.**
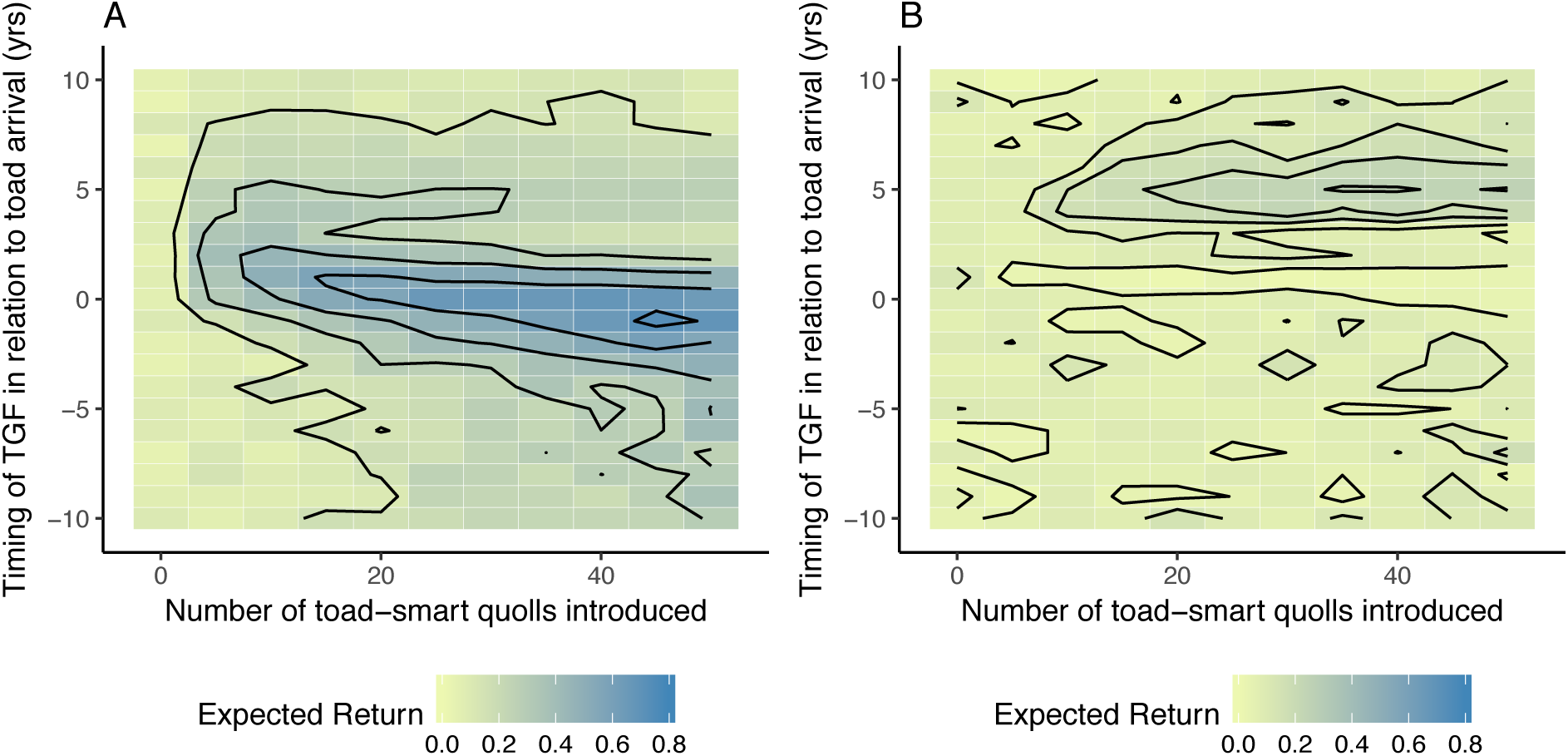
Expected return of the recipient genome (calculated by E(*Y*) = *r*(1 – *p_e_*) using probability of extinction (*p_e_*) and proportion of recipient genome (*r_I_*)) for targeted gene flow with 10% (A) and 50% (B) reduction of fitness for F1 hybrids.

## Discussion

Our model demonstrates that our management objective, *Y,* is sensitive to the timing and size of the introduction. There is also an apparent trade-off between maintaining the local recipient genome, and population survival. Generally, a larger number of introductees in the years immediately preceding and following the stepwise threat produce the lowest probability of extinction, but this also produces lower proportions of recipient genome. The trade-off can be optimised, however, with an introduction of a large number of individuals prior to the arrival of the threat creating the highest expected return for conservation. This strategy produces hybrids in the years prior to toad-arrival so that when selection begins, individuals who carry the recipient genome and toad-smart genes survive. These more optimal strategies retained almost 100% of the recipient population alleles spread across the genome. Our results fit with previous assessments of assisted colonisation that show the timing and number of introductees to be primary considerations for conservation managers undertaking such endeavours (McDonald-Madden *et al.* 2011).

There is, of course, a risk when hybridising populations: outbreeding depression can reduce population fitness (Edmands 2007; Frankham *et al.* 2011). Reduced fitness in hybrids could arise from breakdown of local adaptation, or from genetic incompatibilities, both of which are difficult to predict (Frankham *et al.* 2011). Our model incorporated the possibility of genetic incompatibilities, exploring fitness costs of 10% and 50% accruing to F1 hybrids, and showed outbreeding depression generally had a negative impact on the success of targeted gene flow. These results are unsurprising: we have introduced both a barrier to introgression and a mechanism for reducing fitness. Importantly, however, even with high hybrid dysfunction it was often still beneficial to act. Outbreeding depression did not increase extinction probability above the baseline “do nothing” level, of 95% (scenario where 0 individuals are introduced). Although every situation has its own peculiarities, recent reviews suggest that the risk of outbreeding depression is overstated in the literature: in most realistic cases outbreeding should cause only minor and transitory effects (Aitken & Whitlock 2013). In our particular case, it is likely that northern quolls will experience only minor, if any, effects of outbreeding depression (due to low genetic divergence across their range(Firestone 2000)). The likely fitness benefits gained from carrying favourable alleles that help individuals survive a current and overwhelming threat will likely outweigh any small impact of outbreeding depression (How *et al.* 2009; Weeks *et al.* 2011, 2016), and recombination ensures that maladaptive genetic combinations are rapidly lost.

The scenarios we explored here were limited by the constraints of what is possible for managers of the northern quoll. For example, we limited the number of introductees to what is likely to be logistically possible and used feasible timing and population sizes. The model lacks complexity around genetic architecture: in reality, genes influence traits to varying degrees, loci are varyingly linked, and there are interactions within and between loci (Gomulkiewicz & Holt 1995). For example, dominance in the target trait (which is hinted at in northern quolls(Kelly & Phillips 2018)) would result in a faster adaptive shift. But in the absence of detail on the genetic architecture of this key trait, we have chosen the simplest model possible. In addition, the model was simplified to not include spatial effects, giving all individuals (including introductess) equal chances of survival and finding a mate. In reality, we would expect an introduction to result in some fatalities, so in reality the numbers we are reporting more accurately refer to the number of *surviving* individuals introduced. We also assume that the recipient population is stable prior to toads which, if it is not, may influence the success of the action (McDonald-Madden *et al.* 2011).

Our case study deals with a threat that constitutes a step change in the environment, but there are other threatening processes, such as climate change, that cause gradual change (Kelly & Phillips 2016). Our method of optimising the proportion of recipient genome, *r_I_*, can of course be applied in that context also. For now, however, our results suggest that while timing is important, the optimal timing will depend on the degree of outbreeding depression. Introducing beneficial traits early and generating hybrids prior to the step change is our optimal action, but it becomes less optimal as outbreeding increases, eventually switching to a strategy in which the introduction is best staged after the step change (Fig. 2 & 3). Unfortunately we will rarely know in advance the strength of outbreeding depression. Such uncertainty is common in conservation management (Kujala *et al.* 2013), and modelling, along with decision-theoretic approaches can provide useful framework for optimising actions in particular cases (Regan *et al.* 2005; Polasky *et al.* 2011).

Overall our model suggests that targeted gene flow could provide substantial benefits to populations at risk from an invasive species. No scenario we explored caused more damage than not acting at all. Our model suggests it is possible to use targeted gene flow to reduce population extinction risk while still maintaining the local genetic diversity of the population. We have also identified a useful objective to optimise: the proportion of the recipient population’s genome surviving. This objective directly addresses the trade-off between reducing extinction probability and retaining the recipient genome, and it proved very sensitive to management levers. It would be straightforward, in many circumstances, to anchor this optimisation process with a cost model that adds economic constraints also (e.g., Southwell *et al.* 2017). While each case will have its own particularities, it appears that targeted gene flow could be a valuable tool in an era of rapid environmental change.

## Acknowledgements

This work was supported by the Australian Research Council (LP150100722; FT160100198); Margaret Middleton Fund Award for Endangered Australian Native Vertebrate Animals; and Holsworth Wildlife Research Endowment.

## Supplementary material

### Tables

**Table S1.**
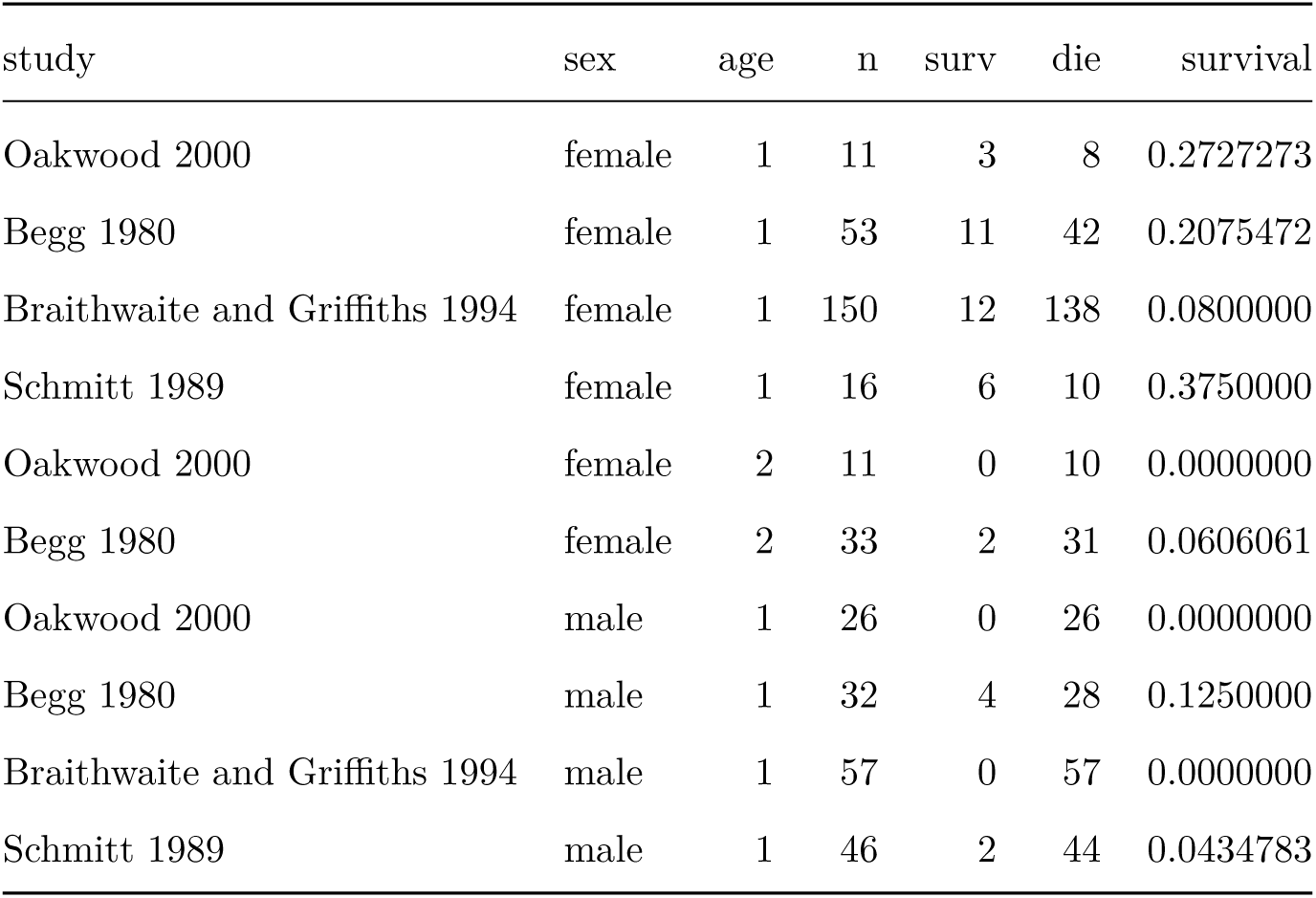
Raw survival data from previous mark-recapture studies used to estimate annual sex-based survival rates

### Heritability sensitivity analysis

We set the heritability of the genes (*h*^2^) to 0.2, which was within the range of behavioural trait heritability given by Roff(Roff 2012). Roff specified that behavioural traits could have a heritability of up to 0.3 however, so we conducted a sensitivity analysis to determine the impact of *h*^2^ on population survival. As would be expected, increased heritability lead to an increase in population survival, and vice versa (Fig. S3 & S4). The selection of *h*^2^ = 0.2 was used in combination of setting other traits (i.e. 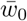, *s*_0_) to initialise a population that acted similar to that of actual quoll populations (95% extinction risk when toads arrived). However it is important to note that heritability plays an important role in the effectiveness of targeted gene flow - but there are still some management techniques that are effective even under very low heritability.

**Figure S1.**
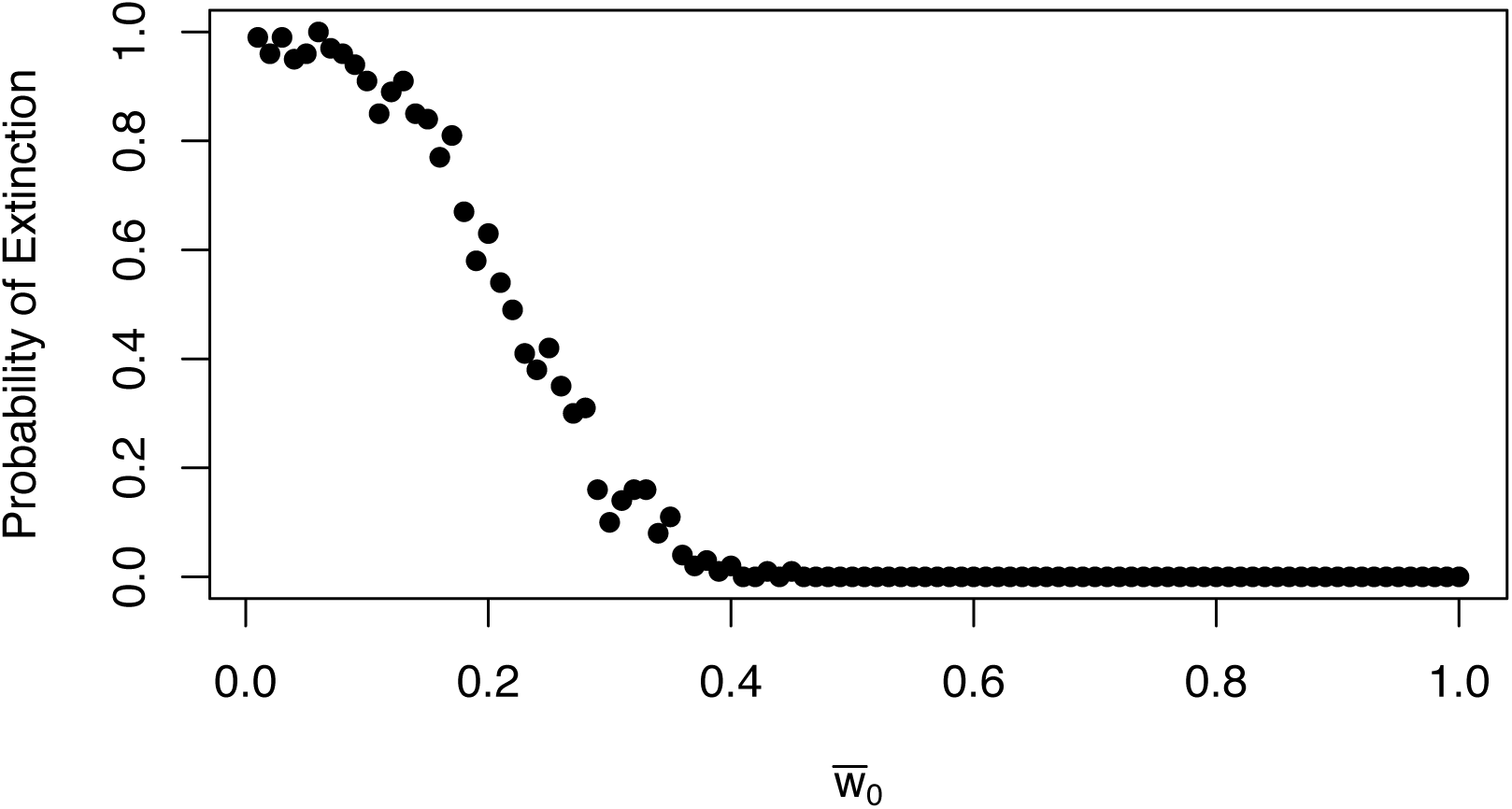
Probability of Extinction of northern quoll population (without targeted gene flow) for values of 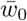 between 0-1.

**Figure S2.**
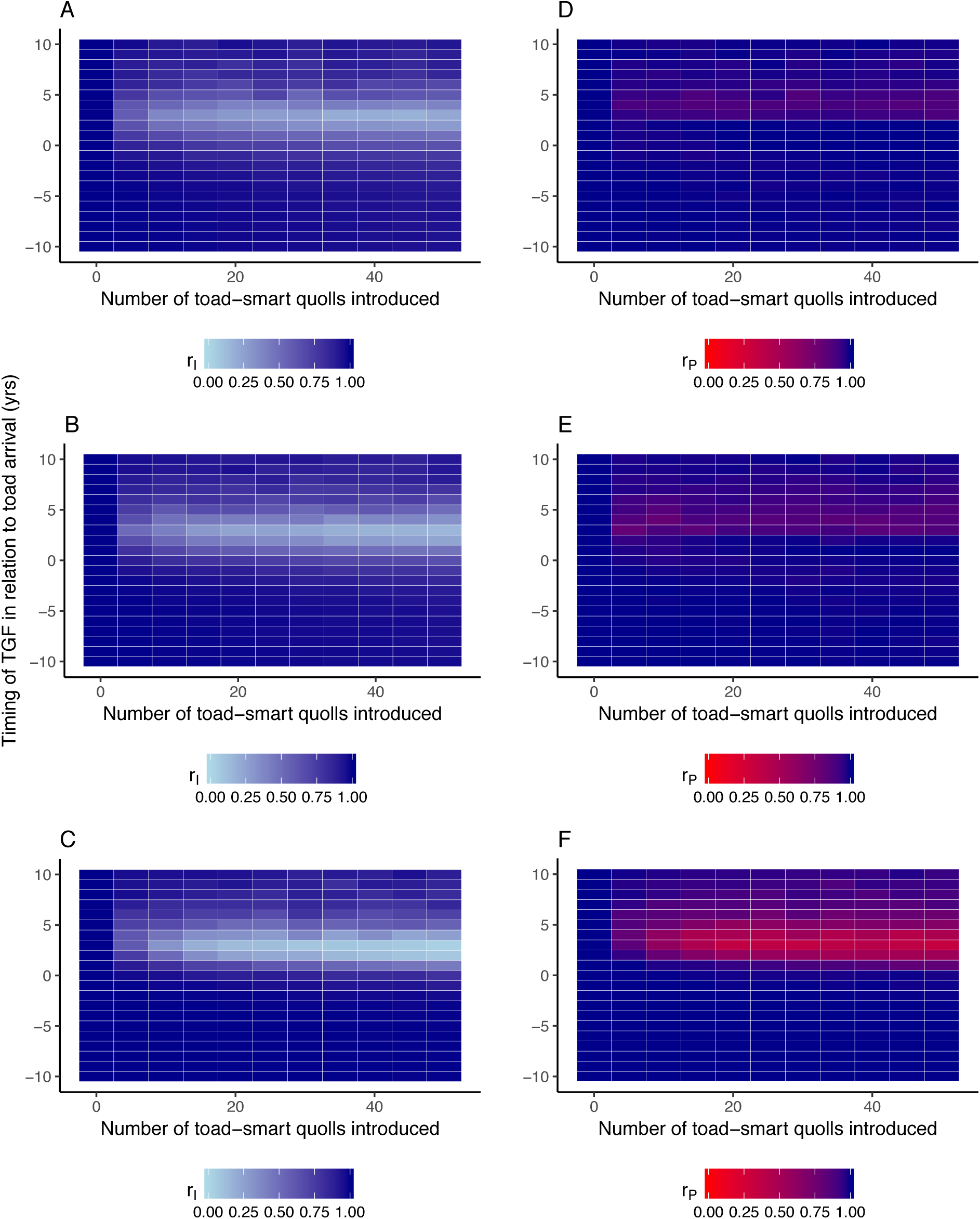
Comparison of two methods for measuring the proportion of recipient genome remaining, *r_I_* and *r_P_.* A-C: *r_I_* calculated as the proportion of recipient alleles remaining in each individual averaged over individuals. D-F: *r_P_* proportion of loci with recipient alleles remaining anywhere within the whole population. A & D: Model without outbreeding depression. B & E: 10% reduction in F1 hybrid fitness. C & F: 50% reduction in F1 hybrid fitness.

**Figure S3.**
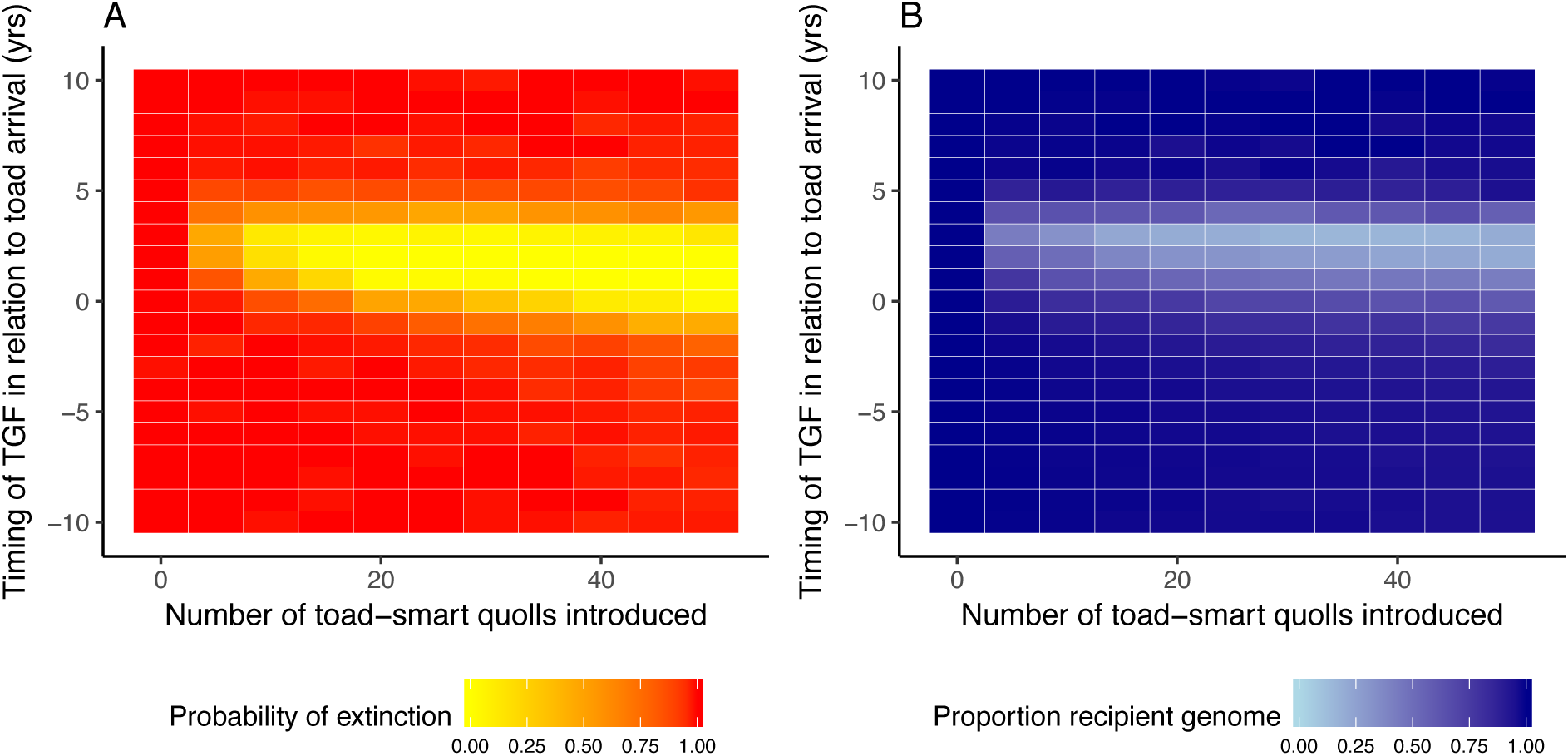
Heritability (*h*^2^) = 0.1. A: The probability of extinction of a northern quoll population (red = high chance of extinction) for varying implementations of targeted gene flow. B: The proportion of recipient population genome (dark blue = recipient orginial genome) in eventual population after varying implementations of targeted gene flow.

**Figure S4.**
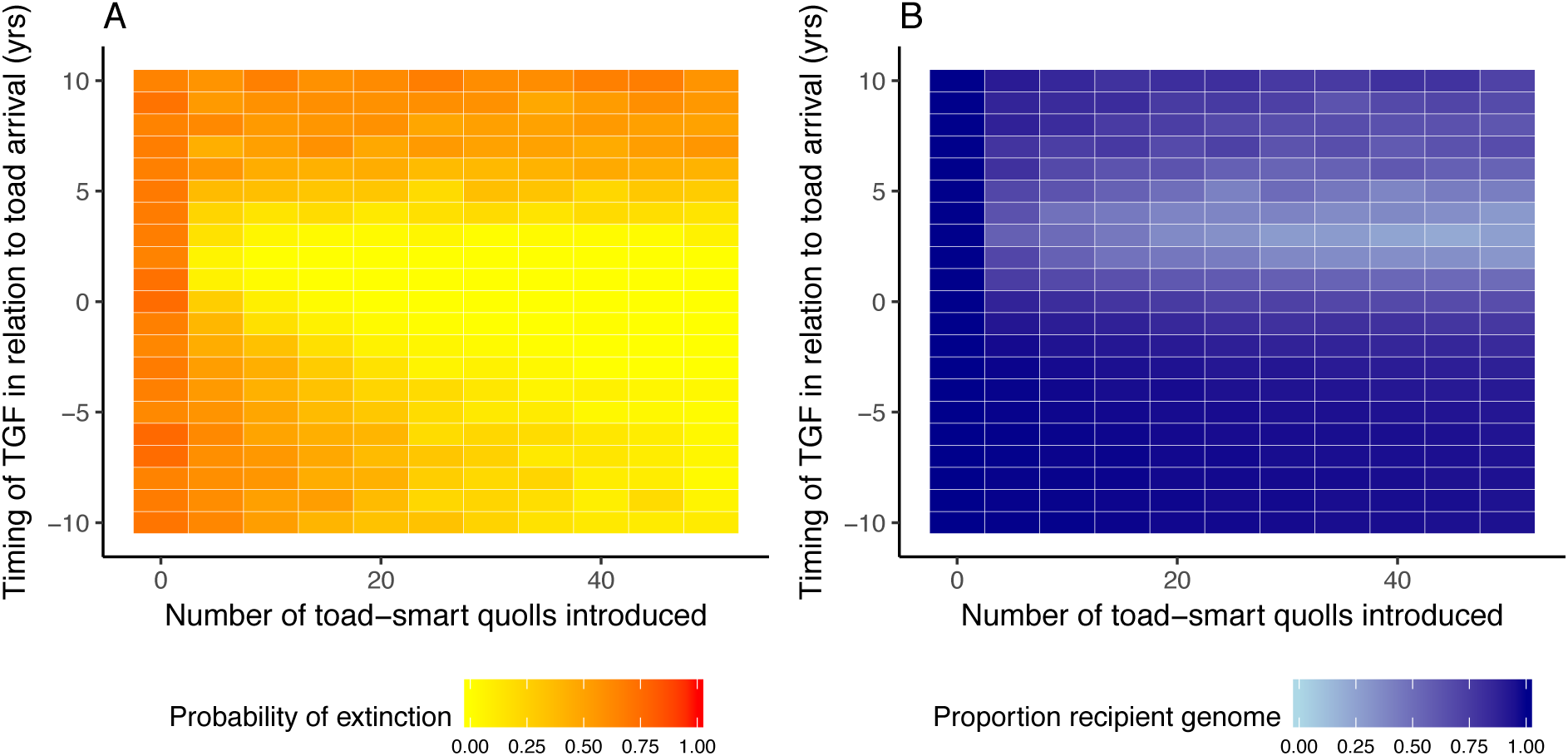
Heritability (*h*^2^) = 0.3. A: The probability of extinction of a northern quoll population (red = high chance of extinction) for varying implementations of targeted gene flow. B: The proportion of recipient population genome (dark blue = recipient orginial genome) in eventual population after varying implementations of targeted gene flow.

